# Genome sequencing of a single tardigrade Hypsibius dujardini individual

**DOI:** 10.1101/053223

**Authors:** Kazuharu Arakawa, Yuki Yoshida, Masaru Tomita

## Abstract

Tardigrades are ubiquitous microscopic animals that play an important role in the study of metazoan phylogeny. Most terrestrial tardigrades can withstand extreme environments by entering an ametabolic desiccated state termed anhydrobiosis. Due to their small size and the non-axenic nature of laboratory cultures, molecular studies of tardigrades are prone to contamination. To minimize the possibility of microbial contaminations and to obtain high-quality genomic information, we have developed an ultra-low input library sequencing protocol to enable the genome sequencing of a single tardigrade *Hypsibius dujardini* individual. Here, we describe the details of our sequencing data and the ultra-low input library preparation methodologies.

## Background & Summary

Tardigrades are small (<1 mm) aquatic Ecdysozoans that occupy a distinct phylum, Tardigrada. This controversial phylogenetic positioning provides important insights into the evolution of arthropods and nematodes^1,2^. Limno-terrestrial tardigrades can withstand almost complete desiccation through a mechanism called anhydrobiosis^3,4^, and tardigrades can survive extreme environments, including extreme temperature^5^, pressure^6,7^, radiation^8–10 11 12^, and even exposure to the vacuum of space. Although it is a weak anhyrobiote, *Hypsibius dujardini* is a model tardigrade due to its ease of culturing^13^, transparent body color that is suited for developmental biology studies, and its feasibility for RNA interference-based genetic studies^14^.

Recently, Boothby *et al*. reported the sequencing of *H. dujardini* and demonstrated that as many as 17.5% of the genes of this organism may have been acquired through horizontal gene transfer (HGT) and that these foreign origin genes may be the key to their extremotolerance^15^. However, the anhydrobiotic capabilities of *H. dujardini* are limited compared with those of other terrestrial tardigrades^12^, and the non-axenic nature of tardigrade culture is prone to contamination, especially in light of the unusually large size of assembly (212.3 Mb) compared with previous estimates^13^. Hence, a calculation of the HGT rate requires comprehensive experimental evidence and careful data screening. Two research groups have provided evidence demonstrating that the HGT candidates presented by Boothby *et al*. were, in fact, mostly contaminant artifacts. Delmont and Eren employed a bioinformatics metagenomics approach to observe the coverage and used a GC-content distribution to identify the contaminants within the assembly.^16^ Koutsovoulos *et al*. presented a thorough comparison using an independently sequenced assembly of the same strain of *H. dujardini* and found no evidence of extensive HGT in this species^17^.

We have also been studying the same strain, but with an ultra-low input sequencing approach. Tardigrades are small animals, mostly in the range of several hundred micrometers in length and comprising approximately one thousand cells, with only several hundred picograms of DNA per individual. Therefore, naive approaches require thousands of animals to extract sufficient DNA or RNA for the construction of libraries for high-throughput sequencing; the collection of that many animals from a non-axenic culture is inherently prone to contamination. Such contamination can be screened after assembly using bioinformatics and metagenomic approaches but would ideally be minimized in the library preparation steps. To this end, we employed an ultra-low input methodology to sequence a single individual of *H. dujardini* to minimize contamination and provided supporting evidence that there is no extensive HGT^18^. Here, we describe the details of our sequencing data and the ultra-low input library preparation methodologies.

## Methods

### Tardigrade culture and sampling

The tardigrade *Hypsibius dujardini* Z151 was purchased from Sciento (Manchester, UK) in August 2012, and its culture was maintained in the laboratory using the culture method previously described for *Ramazzottius varieornatus*^19^, with slight modifications. Briefly, tardigrades were fed *Chlorella vulgaris* (Chlorella Industry) on 2% Bacto Agar (Difco) plates prepared with Volvic water, and the plates were incubated at 18°C under constant dark conditions. Culture plates were renewed every 7~8 days. Adults with body lengths >300 u.m were selected. Cultures were manually inspected every day to obtain freshly laid eggs within 24 h. Eggs were transferred to new plates for egg samples. Similarly, newly laid eggs were observed every day to obtain newly hatched juvenile samples. Anhydrobiotic samples were prepared by placing adult samples in a chamber maintained at 85% relative humidity for 48 h. Successful anhydrobiosis was assumed when >90% of the samples prepared in the same chamber recovered after rehydration.

### Genome sequencing

An adult individual was selected and placed in a sterile agar plate with 1% penicillin streptomycin (Invitrogen) prepared with autoclaved Volvic water for 48 h without food to clear the gut content and remove surface microbes. After 48 h, the animal was repeatedly washed with Milli-Q water on a sterile nylon mesh with 30 u.m pores (Millipore), and any surface microbes were observed at x500~x1000 magnification under a microscope (VHX-5000, Keyence). The cleaned animal was transferred to a low-binding PCR tube with minimal water carry-over (<2 u.1), to which a 200 u.1 genomic lysis buffer (Quick-gDNA MicroPrep kit, Zymo Research) with 0.5% beta-mercaptoethanol was added immediately. The animal was lysed using two freeze-thaw cycles of -80°C and 37°C incubation, and genomic DNA was extracted following manufacturer’s protocol. Extracted DNA was then sheared to 550 bp target fragments with Covaris M220 using a 15 u.1 microTube, and the Illumina library was subsequently prepared using a Thruplex DNA-Seq kit (Rubicon Genomics) according to the manufacturer’s instructions. The purified library was quantified using a Qubit Fluorometer (Life Technologies), and the size distribution was checked using TapeStation D1000 ScreenTape (Agilent Technologies). Libraries with sizes between 400 bp and 1000 bp were selected by manually cutting them out of the agarose and purifying with a NucleoSpin Gel and PCR Clean-up kit (Clontech). The samples were then sequenced using MiSeq v.3 600 cycles kit (Illumina) as 300 bp paired ends. Adapter sequences were removed using the MiSeq software (Illumina).

### mRNA sequencing

Active adults, adults in anhydrobiosis, eggs (1, 2, 3, 4, and 5 days after laying), and juveniles (1, 2, 3, 4, and 5 after hatching), were collected from the culture in three replicates and thoroughly washed with Milli-Q water on a sterile nylon mesh (Millipore). Each individual was placed in a low-binding PCR tube with minimal water carry-over (<2 u.1). Pipette tips were used to crush animals by pressing them against the tube wall, and the individuals were then immediately lysed in TRIzol reagent (Life Technologies). RNA was extracted using a Direct-zol RNA kit (Zymo Research) following the manufacturer’s instructions. RNA quality was checked using High Sensitivity RNA ScreenTape on TapeStation (Agilent Technologies), and the mRNA was amplified via the SMART-Seq approach^20^ using the SMARTer Ultra Low Input RNA Kit for Sequencing v.3 (Clontech). Illumina libraries were prepared using a KAPA HyperPlus Kit (KAPA Biosystems). A purified library was quantified using a Qubit Fluorometer (Life Technologies), and the size distribution was checked using TapeStation D1000 ScreenTape (Agilent Technologies). Libraries with sizes above 200 bp were selected by manually cutting them out of the agarose and purifying with a NucleoSpin Gel and PCR Clean-up kit (Clontech). The samples were then sequenced using a NextSeq 500 High Output Mode 75 cycles kit (Illumina) as single ends. Only two replicates were sequenced for the juvenile day 5 sample. Adapter sequences were removed, and sequences were demultiplexed using the bcl2fastq v.2 software (Illumina).

### Assembly and mapping of sequenced reads for technical validation

Genome sequence reads were assembled *de novo* using MaSuRCA 3.1.3^21^ with a mean insert length of 350 and a standard deviation of 150. Using the assembly and genomic reads, the assembly of Boothby *et al*. (tg.genome.fsa, hereafter referred to as the UNC assembly), one pair of the Boothby reads (TG-500- SIPE_l_sequence.txt and TG-500- SIPE_2_sequence.txt), the assembly of Koutsovoulos *et al*. (nHd.2.3.abv500.fna, hereafter referred to as the Edinburgh assembly) and one pair of the Koutsovoulos reads (gHypdu_hiseq_pe_110427_3_l.fq and gHypdu_hiseq_pe_110427_3_2.fq) were mapped to assemblies in all combinations to assess the mapping rate. Mapping was performed using BWA v.0.7.11 MEM^22^ with the default parameters, and the mapping percentage was calculated using the QualiMap build 11-11-13.

For the RNA-seq data, gene expression abundances were computed using the Kallisto software v.0.42.4 ^23^ with accompanying sleuth utility using the Augustus-based gene prediction data of the Edinburgh assembly (nHd.2.3.1.aug.transcripts.fasta), with the following parameters: -b 100 -bias -single -I 400 -s 50. Overall gene expression profiles were scaled with scale() function and visualized as a heatmap for genes with TPM > 1 using hierarchical clustering based on the Spearman correlation with hclustf) and plotting using the ggplot2 utility in R^24^.

## Data Records

Thirty-six raw Illumina reads were deposited into DDBJ under the BioProject ID PRJDB4575 for one set of whole genome shotgun reads and thity-five sets of RNA-Seq reads comprised of three replicates each (two for the day 5 juveniles) for eggs collected on days 1, 2, 3, 4, and 5 after laying and juveniles collected on days 1, 2, 3, 4, and 5 after hatching (Data Citation 1; see the associated Metadata Record in Table 1 for details). Assembled contigs from the genomic reads were deposited in Data Dryad (Data Citation 2), where the total scaffold length was 132,494,968 bp, the number of contigs was 54,960, the longest contig was 86,209 bp and N50 was 4,851.

**Table 1.** Details of sequenced data.

| BioSample ID | Sample State | Replicate | Type | Data |
| --- | --- | --- | --- | --- |
| SAMD00047018 | Active Adult | 1 | Whole Genome Shotgun | DRR055040 |
| SAMD00047019 | Egg 1 day | 1 | RNA-Seq | DRR055005 |
| SAMD00047019 | Egg 1 day | 2 | RNA-Seq | DRR055006 |
| SAMD00047019 | Egg 1 day | 3 | RNA-Seq | DRR055007 |
| SAMD00047020 | Egg 2 day | 1 | RNA-Seq | DRR055008 |
| SAMD00047020 | Egg 2 day | 2 | RNA-Seq | DRR055009 |
| SAMD00047020 | Egg 2 day | 3 | RNA-Seq | DRR055010 |
| SAMD00047021 | Egg 3 day | 1 | RNA-Seq | DRR055011 |
| SAMD00047021 | Egg 3 day | 2 | RNA-Seq | DRR055012 |
| SAMD00047021 | Egg 3 day | 3 | RNA-Seq | DRR055013 |
| SAMD00047022 | Egg 4 day | 1 | RNA-Seq | DRR055014 |
| SAMD00047022 | Egg 4 day | 2 | RNA-Seq | DRR055015 |
| SAMD00047022 | Egg 4 day | 3 | RNA-Seq | DRR055016 |
| SAMD00047023 | Egg 5 day | 1 | RNA-Seq | DRR055017 |
| SAMD00047023 | Egg 5 day | 2 | RNA-Seq | DRR055018 |
| SAMD00047023 | Egg 5 day | 3 | RNA-Seq | DRR055019 |
| SAMD00047024 | Juvenile 1 day | 1 | RNA-Seq | DRR055020 |
| SAMD00047024 | Juvenile 1 day | 2 | RNA-Seq | DRR055021 |
| SAMD00047024 | Juvenile 1 day | 3 | RNA-Seq | DRR055022 |
| SAMD00047025 | Juvenile 2 day | 1 | RNA-Seq | DRR055023 |
| SAMD00047025 | Juvenile 2 day | 2 | RNA-Seq | DRR055024 |
| SAMD00047025 | Juvenile 2 day | 3 | RNA-Seq | DRR055025 |
| SAMD00047026 | Juvenile 3 day | 1 | RNA-Seq | DRR055026 |
| SAMD00047026 | Juvenile 3 day | 2 | RNA-Seq | DRR055027 |
| SAMD00047026 | Juvenile 3 day | 3 | RNA-Seq | DRR055028 |
| SAMD00047027 | Juvenile 4 day | 1 | RNA-Seq | DRR055029 |
| SAMD00047027 | Juvenile 4 day | 2 | RNA-Seq | DRR055030 |
| SAMD00047027 | Juvenile 4 day | 3 | RNA-Seq | DRR055031 |
| SAMD00047028 | Juvenile 5 day | 1 | RNA-Seq | DRR055032 |
| SAMD00047028 | Juvenile 5 day | 2 | RNA-Seq | DRR055033 |
| SAMD00047029 | Active Adult | 1 | RNA-Seq | DRR055034 |
| SAMD00047029 | Active Adult | 2 | RNA-Seq | DRR055035 |
| SAMD00047029 | Active Adult | 3 | RNA-Seq | DRR055036 |
| SAMD00047029 | Tun Adult | 1 | RNA-Seq | DRR055037 |
| SAMD00047030 | Tun Adult | 2 | RNA-Seq | DRR055038 |
| SAMD00047030 | Tun Adult | 3 | RNA-Seq | DRR055039 |

## Data Citations

1. Arakawa, K., Yoshida, Y., Tornita, M. DDBJ Sequence Read Archive (DRA) DRA004455 (2016).
2. Arakawa, K., Yoshida, Y., Tornita, M. Data Dryad doi:10.5061/dryad.t4k7h (2016).

## Technical Validation

To validate the comprehensiveness of the genomic coverage and to detect any contamination of our single animal protocol, reads and assembled sequences of the three independent projects (UNC, Edinburgh, and the current study) were compared by mapping all possible combinations (Fig. 1). Overall, 84.27% of UNC reads mapped to Edinburgh assembly, which represented reads from *H. dujardini*. The remaining sequences (approximately 10~15%) were likely a contamination of non-tardigrade sequences. A slightly higher percentage (84.72%) of reads mapped to the assembly described in this study, which implied a comparable or slightly greater genome coverage. In addition, 73.95% of the Edinburgh reads mapped to the UNC assembly, whereas a higher percentage (77.10%) mapped to our assembly. Thus, the reads described in this work obtained from a single animal using the ultra-low input protocol yielded genomic reads that were more comprehensive than the traditional methods, which start with thousands of tardigrades. The percentage of mapped reads was lower in the Edinburgh and UNC reads, with sequences likely to be derived from tardigrades in the range of 70~85%, with a contamination in the range of 10~25%. The Edinburgh reads seemed to contain a higher percentage of contamination than the UNC reads. These artifacts were successfully removed during the assembly steps, leaving only 149 contigs spanning 1.3 Mbp. This value represents approximately 1% of the total assembly length, with a coverage less than 1 when using the reads described in this work. By contrast, the UNC assembly contained 5,665 contigs, spanning 56.8 Mbp with a coverage less than 1, which amounted to 22% of the total assembly length. This result indicated that a large proportion of contaminations were contained in the final assembly. By contrast, our reads predominantly mapped to the existing assemblies (94.09% to the UNC assembly, and 97.54% to the Edinburgh assembly). The high rate of mapped reads indicated that contamination was minimal in our data. This result was also supported by the minimal percentage of contigs with coverage less than 1, which was 1% of the total assembly when either the UNC or Edinburgh reads were mapped. Note that our assembly is the unscreened raw output of MaSuRCA assembler and that this percentage would be much lower after screening and curation.

**Figure 1.**
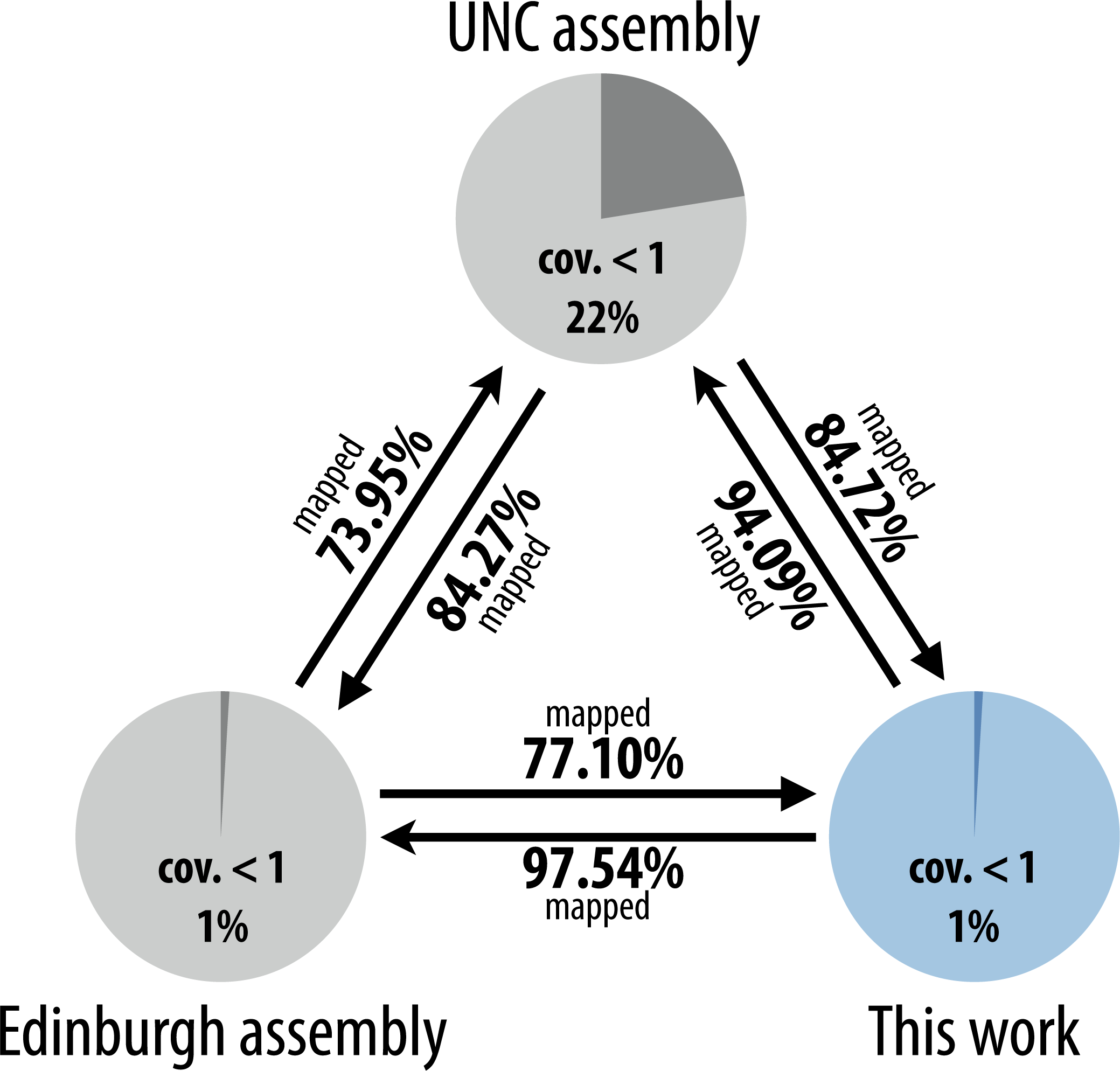
Comparison with existing assemblies and raw reads of *H. dujardini*. The ultra-low input sequencing data described in this work showed greater genomic coverage, as represented by a higher mapping rate of UNC and Edinburgh reads to our assembly, than between that of UNC and Edinburgh. UNC mapped to our assembly at 84.72%, whereas 84.27% mapped to the Edinburgh assembly. Similarly, 77.10% of the Edinburgh reads mapped to our assembly, but only 73.95% mapped to UNC assembly. While the UNC and Edinburgh reads presumably contained 10~25% contamination, as suggested by the percentage of unmapped reads, 94.09% or 97.54% of the study reads mapped to the UNC or Edinburgh assemblies, respectively. These data indicated minimal contamination in our reads.

The mapping percentage and abundance estimates of the RNA-Seq data were visualized for active and tun (in anhydrobiosis) adult samples (Fig. 2) and for juvenile and egg samples (Fig. 3). Adult samples exhibited high mapping percentages and expression profiles, which mainly correlated within the biological replicates. However, both the reproducibility and mapping percentages were consistently low, with the exception of juvenile samples on the first and third days after hatching. This result is presumably due to the inefficiency of RNA extraction in these developmental stages.

**Figure 2.**
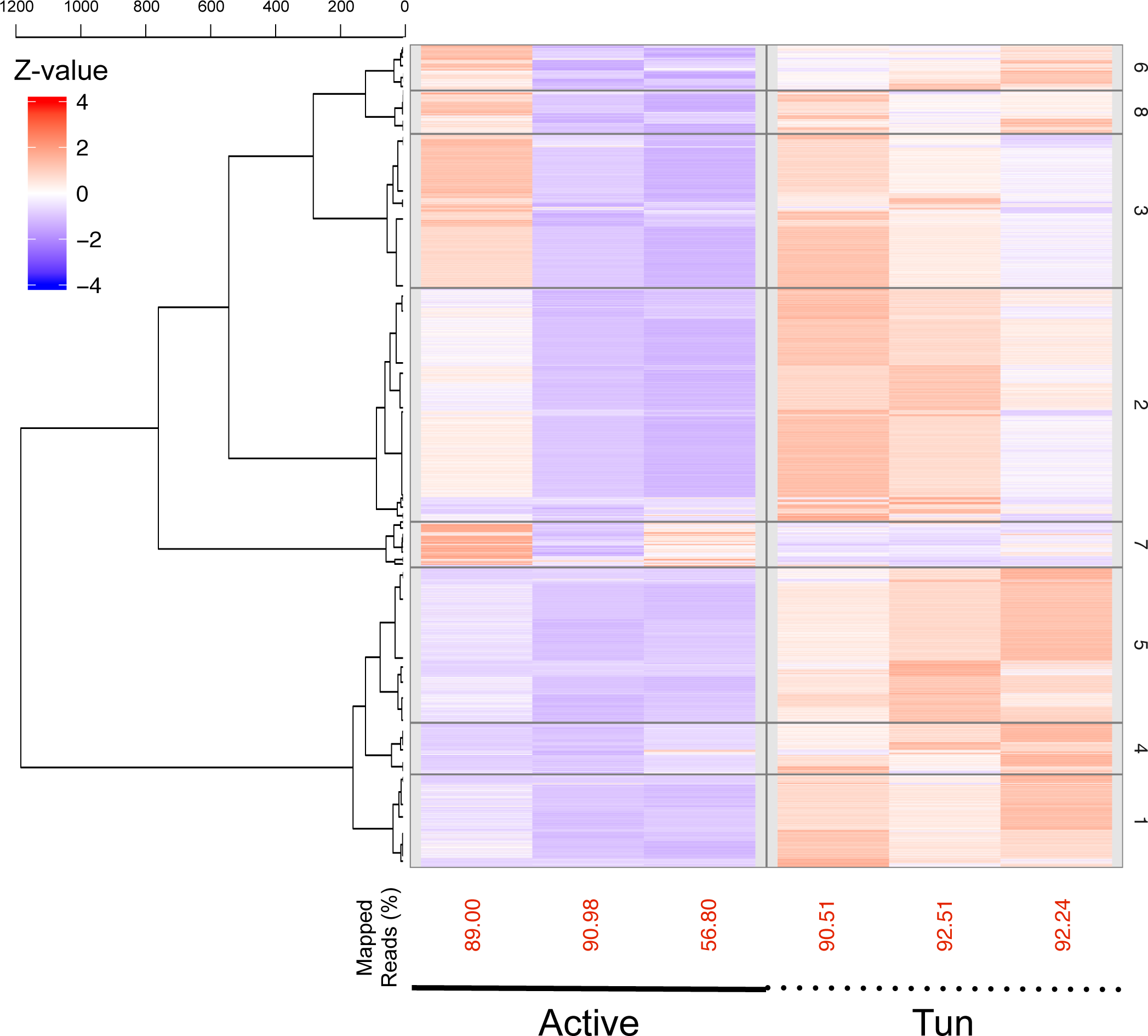
Transcriptome abundances of active and tun (in anhydrobiosis) adults in three biological replicates. Percentage of mapped reads is shown beneath the heatmap.

**Figure 3.**
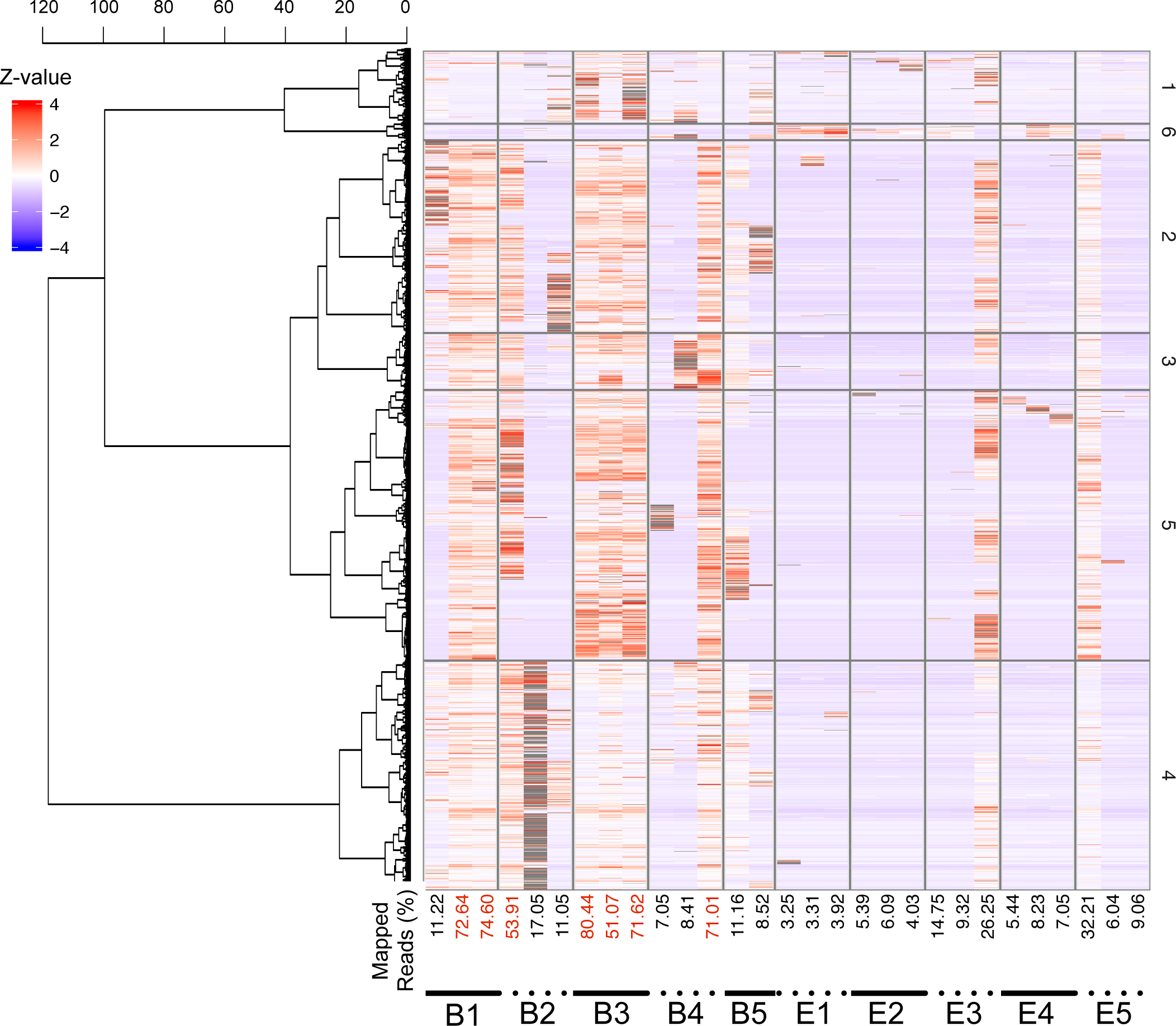
Transcriptome abundances of juveniles (B1~B5) and eggs (E1E5) in the first 1~5 days after hatching or laying. Sequences were obtained in three biological replicates except for B5, which only has two replicates. The percentage of mapped reads is shown beneath the heatmap, and samples with a mapping percentage greater than 50% are colored in red.

## Usage Notes

The genome sequencing reads described in this work were intended to confirm the published UNC and Edinburgh assemblies and to clarify the contigs derived from *H. dujardini*, as described previously^18^. Moreover, the assembly provided in this work was intended only for technical validation, as described above. The assembly was neither scaffolded nor curated. Therefore, the Edinburgh assembly would be better suited for genomic studies. The RNA-Seq data described in this work were obtained to further validate the confirmed data. As described in the technical validation section, the reproducibility was low for samples in the early developmental stages, presumably due to inefficient RNA extraction from eggs and juveniles. RNA-Seq data with low mapping percentage should not be used in comparative analyses.

## Acknowledgments

The authors thank Nozomi Abe for technical support in tardigrade culturing and genomic sequencing, and Yuki Takai for technical support in RNA sequencing. *Chlorella vulgaris* used to feed the tardigrades was provided courtesy of Chlorella Industry Co. LTD. This work was supported by KAKENHI Grant-in-Aid for Young Scientists (No.22681029) from the Japan Society for the Promotion of Science (JSPS), by a Grant for Basic Science Research Projects from The Sumitomo Foundation (No. 140340), and partly by research funds from the Yamagata Prefectural Government and Tsuruoka City, Japan.

## Author contributions

KA designed and performed the experiments and drafted the manuscript. YY analyzed the RNA-seq data. MT managed the computer resources. All authors contributed to editing and revising the manuscript.

